# The first genetic linkage map for *Fraxinus pennsylvanica* and syntenic relationships with four related species

**DOI:** 10.1101/365676

**Authors:** Di Wu, Jennifer Koch, Mark Coggeshall, John Carlson

## Abstract

Green ash (*Fraxinus pennsylvanica*) is an outcrossing, diploid (2n=46) hardwood tree species, native to North America. Native ash species in North America are being threatened by the rapid invasion of emerald ash borer (EAB, *Agrilus planipennis*) from Asia. Green ash, the most widely distributed ash species, is severely affected by EAB infestation, yet few resources for genetic studies and improvement of green ash are available. In this study, a total of 5,712 high quality single nucleotide polymorphisms (SNPs) were discovered using a minimum allele frequency of 1% across the entire genome through genotyping-by-sequencing. We also screened hundreds of genomic- and EST-based microsatellite markers (SSRs) from previous *de novo* assemblies (Staton et al. 2015; Lane et al. 2016). A first genetic linkage map of green ash was constructed from 91 individuals in a full-sib family, combining 2,719 SNP and 84 SSR segregating markers among the parental maps. The consensus SNP and SSR map contains a total of 1,201 markers in 23 linkage groups spanning 2008.87cM, at an average inter-marker distance of 1.67 cM with a minimum logarithm of odds (LOD) of 6 and maximum recombination fraction of 0.40. Comparisons of the organization the green ash map with the genomes of asterid species coffee and tomato, and genomes of the rosid species poplar and peach, showed areas of conserved gene order, with overall synteny strongest with coffee.

## Introduction

Green ash (*Fraxinus pennsylvanica* Marsh.) is the most widely distributed angiosperm tree in North America. This species, along with other native ash species, has been threatened by the rapid invasion of the emerald ash borer (EAB) insect from Asia. The economic value of ash trees (*Fraxinus spp*.) is difficult to assess, but as the most widely distributed ash species in NA, green ash may contribute billions of dollars to the economy, especially when used as a preferred urban street tree (Kovacs et al., 2010). Unfortunately, EAB has resulted in the death of hundreds of millions of ash trees (Herms and McCullough, 2014), which cost municipalities, property owners, nursery operators and forest product industries hundreds of millions of dollars. Current studies of ash species have included transcriptome analyses (Lane et al. 2016; Bai et al. 2011; Sollars et al. 2017), metabolite comparisons (Whitehill et al. 2012; Sambles et al. 2017), and a *de novo* genome assembly for European ash, *Fraxinus excelsior*(Sollars et al. 2017). However no genetic linkage maps have yet been reported for ash species.

The genetic and molecular mechanisms underlying diverse biological traits in green ash are unknown. Genetic linkage maps are powerful tools for genomics and genetics research. With a large number of DNA markers, fine mapping of quantitative trait loci (QTL) can support marker-assisted selection for breeding programs. Candidate genes can also be positioned within the quantitative trait locus (QTL) regions identified by association studies. Combined with transcriptome studies, differentially expressed genes can be mapped and localized so that genes related to specific traits of interest can be characterized. In addition, linkage maps can also be used to conduct comparative analyses with other species to detect genomic synteny and to improve and correct chromosome-scale genome assemblies. Linkage maps have been reported in many non-model species (Tian et al. 2015; Moumouni et al. 2015; Raman et al. 2014), but the resolution of genetic maps depends on the number of markers segregating in the mapping population.

Recently, the innovation of NGS methods for genetic marker discovery combined with reduced costs has facilitated the genotyping of thousands of SNPs across hundreds of samples. New methods, such as reduced-representation libraries (RRLs) (Van Tassell et al. 2008), restriction-site associated sequencing (RAD-seq) (Davey and Blaxter 2010) and genotyping-by-sequencing (GBS) (Elshire et al. 2011), utilize restriction enzyme digestion of target genomes to reduce the complexity of the target. In addition, new genotyping approaches including Illumina’s BeadArray™ technology based GoldenGate^®^ and Infinium^®^ assays, and the Sequenom MassARRAY^®^technology (Gabriel, Ziaugra, and Tabbaa 2001), facilitate SNP genotyping at the genome-wide level in a cost-effective manner. However, as biallelic markers, SNPs are usually less informative than multi-allelic makers, such as microsatellites, which can amplify up to four alleles in a diploid species. Therefore, SSRs are still robust markers for use in genetic map development and many other genetic studies, which justifies their use despite how laborious and time-consuming the development and use of microsatellite markers in genotyping can be.

## Materials and Methods

### Plant material and DNA extraction

A controlled cross pollination between an EAB-resistant and an EAB-susceptible parent was performed at the USDA Forest Service, Northern Research Station in Delaware, OH and produced an F1 population of 780 individuals. The maternal parent tree was a grafted ramet of a green ash tree located at the Dawes Arboretum in Newark, OH, and the male (pollen) parent tree was a grafted ramet of a tree located in the Oak Openings Metropark in Toledo, OH, that retained a healthy canopy despite long-term EAB-infestation in this area (Knight et al. 2012). The maternal parent tree (PE0048) was heavily infested by EAB and considered susceptible, while the paternal parent tree (PE00248) is considered to have a level of resistance to EAB, based on its field phenotype and confirmed by replicated EAB-egg inoculation tests performed as described by Koch (Koch et al. 2015). During growth of the seedlings in the greenhouse, an unspecified viral infection reduced the family size to 543 survivors, which were transferred to a gravel-bed nursery at the Center for Agroforestry, University of Missouri, Columbia, MO. For this genetic linkage mapping study, leaf samples were collected from 91 of the seedlings and from the parent trees for genomic DNA extraction. DNA was isolated from frozen leaves using a CTAB method (Clarke 2009). DNA samples were quantified using a Qubit 2.0 fluorimeter (Invitrogen, Carlsbad, CA, USA) and then checked for molecular weight in 0.8% agarose gel. The 91 progeny were verified by SSR analysis as authentic full siblings using Colony parentage analysis (Jones and Wang 2010).

### Genotyping-by-sequencing

GBS libraries were constructed and sequenced at the Cornell University Genomic Diversity Facility, using the protocol described by Elshire (Elshire et al. 2011). Three methylation-sensitive restriction enzymes (i.e. *Ape*KI, *EcoT22I* and *Pst*I) were tested prior to selection of *Pst*1 as the best for GBS library construction. A compatible set of 96 barcode sequences were included in the library construction for subsequent multiplex sequencing. The libraries of 95 samples (91 full-sib progeny and 2 libraries for male and female parents) and a blank negative control were pooled and sequenced using Illumina NextSeq 500 on one lane of single-end reads, running for 86 cycles to obtain a read length of 86bp.

### GBS Data Processing and SNP discovery

The TASSEL-GBS pipeline was used for SNP discovery and SNP calling (Glaubitz et al. 2014), using a draft genome assembly of green ash, consisting 495,002 contigs (Buggs, unpublished) as a reference. The raw data FASTQ files were processed for sequence filtering through the pipeline using a minimum Qscore of 20 across the first 64 bases. Raw reads with a perfect match to one of the barcodes plus the subsequent five nucleotides that were expected to remain from a *PstI* cut-site (i.e. 5’…CTGCA’G…3’) and without N’s were retained as good barcoded reads. Identical good barcoded reads were then clustered into tags, while rare tags represented by fewer than 5 reads were excluded from the dataset. Each tag was then aligned to the reference genome and only genomic positions of the tags that aligned to a unique best position in the genome were kept for further processing. SNP discovery was performed for each set of tags that align to the same starting genomic position. For each SNP, the allele represented by each tag was then determined with the observed depth of each allele. The initial filtering process was based upon minimum allele frequency (MAF) of 0.01 to remove error-prone and spurious SNPs. To exclude less reliable SNPs, only SNPs with less than 10% missing genotyping data across samples were kept for downstream analysis.

### Simple sequence repeats (SSRs) genotyping and data analysis

*De novo* assemblies of the green ash genome and transcriptome have been used previously to search di-, tri-, and tetra-nucleotide microsatellite repeats (Lane et al. 2016). Primers for simple sequence repeats (SSRs) loci were designed using default settings in PRIMER3 (Untergasser et al. 2012; Koressaar and Remm 2007). A total of 419 primer pairs were tested within the mapping population. Total genomic DNA samples were normalized to 2ng/μl and then used for PCR amplification reactions using 5x FIREPol Master Mix (Solis BioDyne) and custom primer mix. To reduce the cost of fluorescent primers, a three-primer system was used, which included a universal M13 oligonucleotide (TGTAAAACGACGGCCAGT) labeled with fluorescent dye, a sequence-specific forward primer with the M13 tail at its 5’ end, and a sequence-specific reverse primer (Schuelke 2000). Conditions of the PCR amplification were as follows: 95°C (15 min), then 35 cycles at 94°C (30 s) / 56°C (90 s) / 72°C (90 s) and a final extension at 72°C for 10 min. Pooled PCR reaction products were sized on an ABI 3730xl capillary electrophoresis instrument, with peaks identified by *GeneScan* (Applied Biosystems) followed by fragment sizing using *GeneMapper* v5.0 (Applied Biosystems).

### Linkage map construction

The allele data for all selected markers were tested for segregation distortion and genotypic data similarities. Markers that showed significantly distorted segregation (P<0.05) were removed from the dataset. Redundant markers whose genotypic data showed 100% similarity were also removed from the dataset to improve the computation efficiency. Male and female-specific linkage maps were then constructed individually for the filtered markers with JoinMap 4.0 using the regression algorithm and Kosambi mapping function and designating a cross-pollinator (CP) population type (Van Ooijen 2006). In the first run, only markers with up to 5% missing data and categorized as lm x ll, nn x np, ef x eg and ab x cd allele combinations were used to build the framework maps (LOD score threshold = 6). For maternal maps, the genotype codes from loci with segregation types <ef x eg> and <ab x cd> were translated to genotype codes of <lm x ll> and for paternal maps to genotype codes of <nn x np>. In the second run, loci with segregation type <hk x hk> and less informative markers with 5% to 10% missing data were included to build the maps, while using the marker order obtained from step one as the Start Order in JoinMap. The genotypes hk, hh and kk were translated to unknowns, ll and lm, respectively for maternal parent and to unknowns, nn and np for paternal parent. The consensus map was then established by integrating both paternal and maternal maps through shared markers using *MergeMap* (Wu, Close, and Lonardi 2008). The linkage maps were then drawn using *MapChart* 2.2 (Voorrips 2002).

### Comparative analysis with genetic maps of two Asterid and two Rosid species

Fifty base pairs of flanking sequences from both sides of the mapped green ash SNP markers were searched against the tomato, poplar and peach genomes (Phytozome v10.0) and coffee genome (Coffee Genome Hub) using BLASTN with an e-value threshold of 1e-5. Circos v0.69 was then used to plot synteny between green ash genetic maps and chromosomes of tomato (Krzywinski et al. 2009), coffee (Denoeud et al. 2014), peach (Arús et al. 2012) and poplar (Tuskan et al. 2006). To plot the syntenic regions, cM distances on the genetic maps of green ash were converted to base pairs using an averaged cM/bp value, based on the total linkage length in cM of the map and the estimated total genome size of green ash (i.e. 961Mb) (Staton et al, 2015).

## Results

### Enzyme selection

We tested whether one of the commonly used restriction enzymes would generate an appropriate distribution of fragment lengths across the genome of green ash (Fig. 1). Fragment size distribution was checked on the Bioanalyzer 2100 HS-DNA chip (Agilent Technologies, Santa Clara, CA, USA) (Fig. 1). All three enzymes yielded a large number of fragments ranging between 150bp and 500bp, which were suitable for the GBS approach. However, since both *ApeKI* and *EcoT22I* showed some repetitive peaks and a small proportion of fragments greater than 500bp, we selected the enzyme *PstI* to construct GBS libraries for green ash.

**Figure 1:**
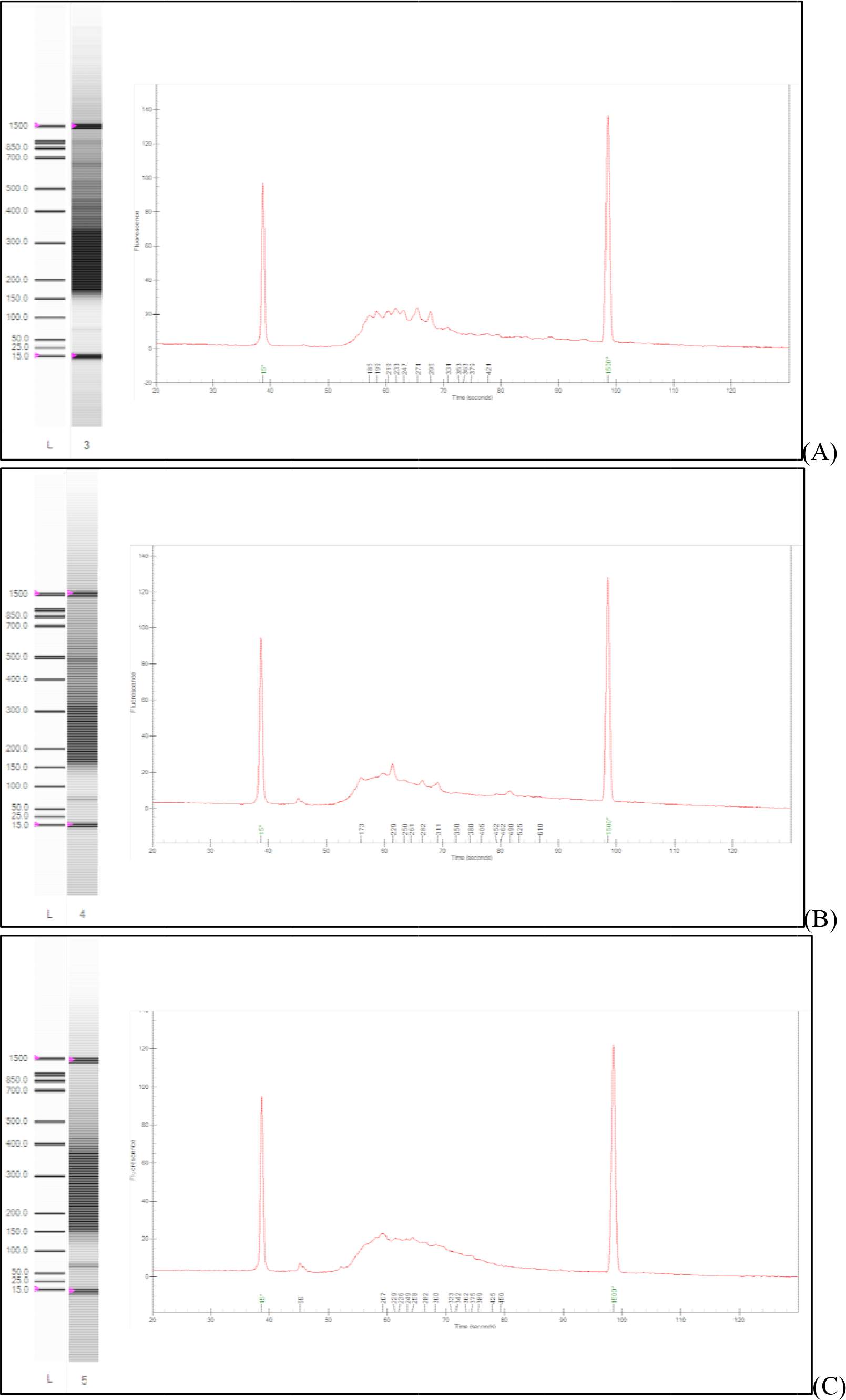
Assessment of GBS libraries using three different restriction enzymes. (A) GBS libraries prepared with *ApeKI*; (B) GBS libraries prepared with *EcoT22I*; (C) GBS libraries prepared with *PstI*.

### Genome-wide identification of SNPs

To identify genome-wide SNPs from green ash, the 6-base cutter restriction enzyme *PstI* was used to digest the genome and construct the 96-plex GBS libraries of the 91 individuals in the F1 population and their male and female parents. A total of 63,540 putative SNPs that had a MAF of 1% were identified using the TASSEL GBS pipeline. With a maximum of 20% missing genotype data, 5,712 SNPs were retained for map construction. The frequency of SNP occurrence across the genome is summarized in Table 1. In general, the frequency of transitions (63.65%) was higher than transversions (35.83%). The most widespread variation was A/G (31.86%) while the least common variation was C/G, accounting for 6.53% of the total detected SNPs. We observed a transition:transversion (Ts/Tv) ratio of 1.78, which was similar to the observations for other plant species (Gaur et al. 2015; Pootakham et al. 2015). A set of bi-allelic SNPs were scored as lm x ll, nn x np and hk x hk. By using a stringent cutoff of 10% missing data, 2,729 high quality and polymorphic SNPs were retained for further analysis. A total of 727 and 1,548 SNPs were polymorphic in maternal and paternal parents, respectively, while 454 SNPs were polymorphic in both parents.

**Table 1.** Summary of SNPs identified in green ash.

|  |  |  |
| --- | --- | --- |
| Total number of SNPs | 5712 | 100% |
| Bi-allelic | 5683 | 99.49% |
| others | 29 | 5.01% |
| Transversion |  |  |
| A/C | 494 | 8.65% |
| A/T | 652 | 11.41% |
| C/G | 373 | 6.53% |
| G/T | 528 | 9.24% |
| Transition |  |  |
| A/G | 1820 | 31.86% |
| C/T | 1816 | 31.79% |

**Table 2.** Summary of SSR markers.

| Type | Tested | Polymorphic | Segregation type |  |  |  |  |
| --- | --- | --- | --- | --- | --- | --- | --- |
|  |  |  | ab x cd | ef x eg | hk xhk | ll x lm | nn x np |
| EST-SSR | 352 | 30 | 14 | 9 | 1 | 5 | 1 |
| gSSR | 252 | 54 | 25 | 12 | 0 | 7 | 10 |

### Polymorphism of EST-derived and genomic SSR markers in F1 mapping population

SSR polymorphism assessment was first examined in six randomly selected F1 progeny and the two parents using non-fluorescent primers. Among the 352 EST-derived and 252 genomic SSR primer pairs, 84 (13.91%) successfully amplified polymorphisms between the two mapping parents, including 30 EST-SSR and 54 gSSR markers. The 84 pairs of primers and related information including primer name, motif type, forward and reverse primers, and expected product size are listed in Table S1. Sequences associated with 30 EST-SSRs were aligned by BLASTX against the GenBank non-redundant (nr) protein database with an e-value cut-off of 1e-5. The BLASTx search results showed that 19 (63.33%) of the 30 polymorphic EST-SSR loci are markers for known or uncharacterized protein coding genes (Table S1).

### Construction of genetic linkage maps

Chi-square tests were performed to test for deviation from the expected Mendelian segregation ratio. Of the 2731 polymorphic SNPs (<10% missing data), 902 markers (33.02%) showed significant segregation distortion (P ≤ 0.05). Severe distortion (SD) was detected in 604 SNP markers (P≤0.001) while 298 SNP loci were moderately distorted (0.001<P≤0.05).

After removing the markers that significantly deviated from the expected Mendelian ratio (P≤0.05), a total of 90 samples with genotypic data for 2049 SNP and SSR loci were retained for genetic map construction. To reduce computation time, loci with identical genotypes were eliminated. A set of 1537 high quality SNP loci along with 75 SSR markers that segregated in 90 members of the F_1_ population arising from the intra-specific cross of PE00248 and PE0048 were used to construct an intra-specific linkage map for *F. pennsylvanica*.

The female genetic map consisted of 992 markers mapped on 760 distinct positions spanning 1562.64cM, with an average marker interval of 2.22cM, ranging from 1.08cM in LG9 to 3.97cM in LG22. Among the 992 markers, 6 markers were later assigned individually to LG21, but it was not possible to compute genetic distance between the markers (Table 3) and thus, a linkage group could not be generated. As a result, only 22 LGs were constructed in the maternal map. The male genetic map consisted of 755 markers on 1744.12cM, with an average marker interval of 2.87cM (Table 3), ranging from an average paired marker distance of 1.16 cM in LG18 to 4.99 cM in LG21. In total, 230 markers were shared between the two parental maps, which were used to construct the consensus map by integrating the two parental maps at those loci. Overall, the average male-to-female ratio of map length was 1.12, ranging from 0.36 in LG18 to 2.3 in LG16.

**Table 3.** Summary of female and male genetic maps.

| Female (PE0048) |  |  |  |  | Male (PE00248) |  |  |  |  | Common markers | M:F |
| --- | --- | --- | --- | --- | --- | --- | --- | --- | --- | --- | --- |
| LGs | Markers | Distinct positions | Length | Average interval | LGs | Markers | Distinct positions | Length | Average interval |  |  |
| 1 | 71 | 55 | 97.69 | 1.78 | 1 | 46 | 39 | 115.76 | 2.97 | 2 | 1.18 |
| 2 | 33 | 28 | 66.65 | 2.38 | 2 | 49 | 42 | 99.14 | 2.36 | 10 | 1.49 |
| 3 | 54 | 39 | 76.49 | 1.96 | 3 | 38 | 33 | 102.38 | 3.1 | 14 | 1.34 |
| 4 | 48 | 40 | 92.16 | 2.3 | 4A | 22 | 16 | 48.51 | 3.03 | 9 | 0.7 |
|  |  |  |  |  | 4B | 5 | 4 | 15.66 | 3.92 | 4 |  |
| 5 | 54 | 43 | 103.21 | 2.4 | 5 | 33 | 27 | 75.71 | 2.8 | 11 | 0.73 |
| 6 | 43 | 35 | 80.91 | 2.31 | 6 | 33 | 29 | 92.78 | 3.2 | 13 | 1.15 |
| 7 | 51 | 40 | 75.62 | 1.89 | 7 | 42 | 34 | 83.77 | 2.46 | 14 | 1.11 |
| 8 | 49 | 39 | 78.3 | 2.01 | 8 | 34 | 30 | 91.48 | 3.05 | 10 | 1.17 |
| 9 | 97 | 69 | 74.72 | 1.08 | 9 | 49 | 48 | 85.83 | 1.79 | 23 | 1.15 |
| 10 | 42 | 31 | 74.96 | 2.42 | 10 | 41 | 32 | 90.52 | 2.83 | 12 | 1.21 |
| 11 | 30 | 25 | 64.96 | 2.6 | 11 | 36 | 31 | 90.58 | 2.92 | 5 | 1.39 |
| 12 | 43 | 36 | 77.53 | 2.15 | 12A | 28 | 18 | 71.01 | 3.95 | 10 | 1.03 |
|  |  |  |  |  | 12B | 6 | 5 | 8.63 | 1.73 | 1 |  |
| 13 | 45 | 33 | 67.47 | 2.04 | 13 | 39 | 37 | 80.29 | 2.17 | 13 | 1.19 |
| 14 | 37 | 29 | 72.15 | 2.49 | 14 | 32 | 27 | 78.59 | 2.91 | 10 | 1.09 |
| 15 | 36 | 27 | 46.06 | 1.71 | 15 | 25 | 20 | 85.34 | 4.27 | 5 | 1.85 |
| 16 | 19 | 15 | 32.29 | 2.15 | 16 | 42 | 35 | 74.27 | 2.12 | 7 | 2.3 |
| 17 | 59 | 47 | 67.51 | 1.44 | 17 | 25 | 25 | 62.58 | 2.5 | 10 | 0.93 |
| 18 | 49 | 37 | 68.47 | 1.85 | 18 | 26 | 21 | 24.33 | 1.16 | 13 | 0.36 |
| 19 | 23 | 20 | 66.47 | 3.32 | 19 | 22 | 20 | 62.36 | 3.12 | 8 | 0.94 |
| 20 | 33 | 22 | 67.11 | 3.05 | 20 | 18 | 14 | 49.48 | 3.53 | 7 | 0.74 |
| 21 | 6 | -- | -- | -- | 21 | 17 | 14 | 69.86 | 4.99 | -- | -- |
| 22 | 23 | 15 | 59.56 | 3.97 | 22 | 25 | 18 | 51.55 | 2.86 | 10 | 0.87 |
| 23 | 53 | 35 | 52.35 | 1.5 | 23 | 22 | 16 | 33.71 | 2.11 | 9 | 0.64 |
| Total | 992 | 760 | 1562.6 | 2.22 |  | 755 | 635 | 1744.1 | 2.87 | 230 | 1.12 |

The total length of the consensus map was 2008.98cM with LG1 (125 cM) being the largest and LG23 (53.93cM) being the smallest (Fig. 2, Table 4). In total, 1201markers were placed on the consensus map, consisting of 75 SSR and 1126 SNP markers. The number of markers per linkage group varied from 13 (LG21) to 95 (LG9), with an average of 52 markers per linkage group.

**Figure 2:**
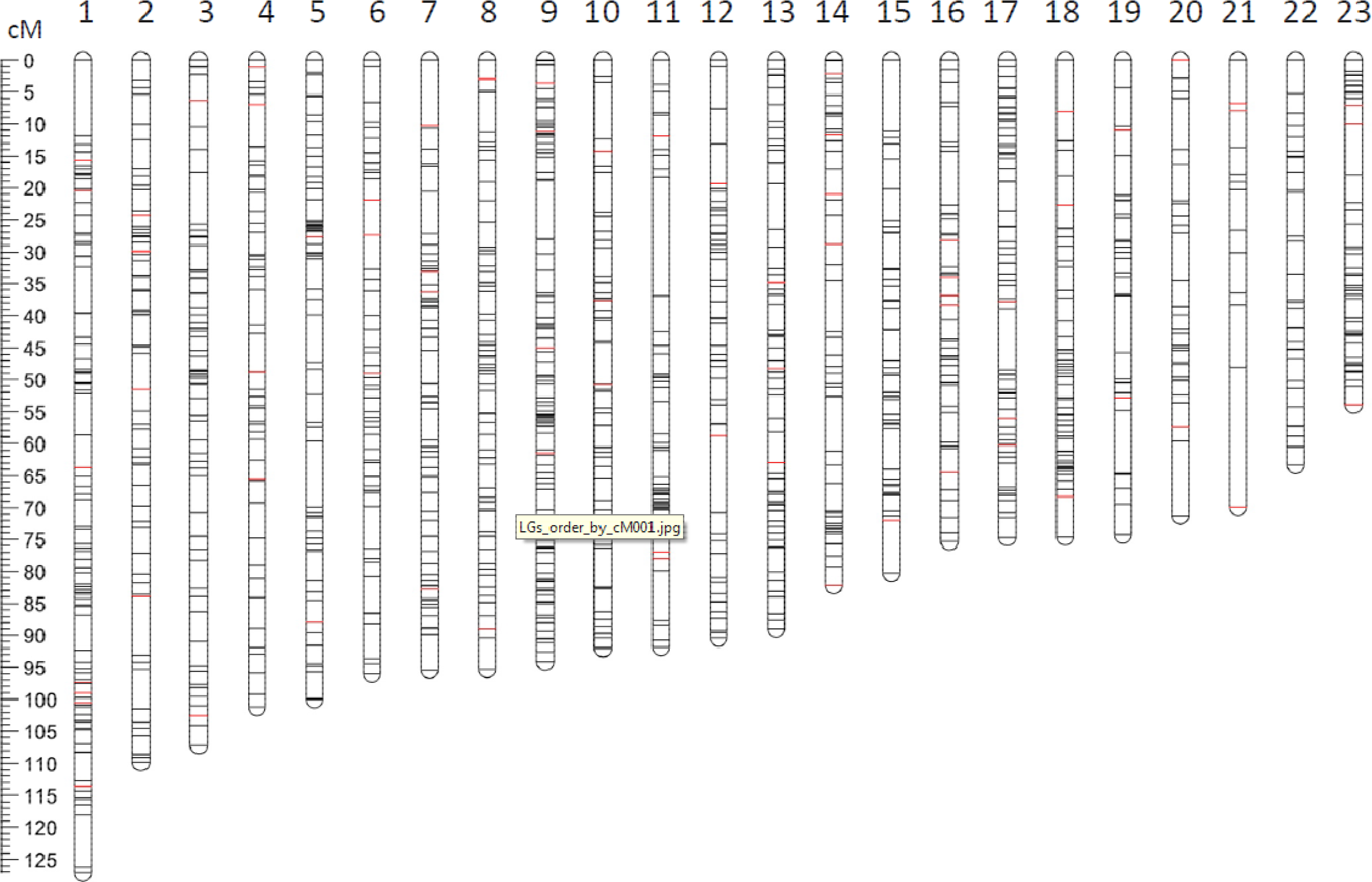
Genetic maps of *F. pennsylvanica*. The intra-specific linkage map of green ash based on F1 population of one EAB-resistant and one EAB-susceptible genotypes harboring 1201 loci. SNP markers are represented in black, while red color represented SSR markers.

**Table 4.** Summary of genetic linkage map of *F. pennsylvanica*.

| LGs | mapped markers |  | Genetic size (cM) | Average marker interval | Max gap (cM) |
| --- | --- | --- | --- | --- | --- |
|  | SSRs | SNPs |  |  |  |
| 1 | 7 | 85 | 127.05 | 1.38 | 11.97 |
| 2 | 5 | 56 | 109.81 | 1.8 | 9.15 |
| 3 | 2 | 59 | 107.22 | 1.76 | 8.06 |
| 4 | 4 | 47 | 101.17 | 1.98 | 6.51 |
| 5 | 2 | 58 | 99.99 | 1.67 | 10.31 |
| 6 | 3 | 48 | 96.01 | 1.85 | 6.61 |
| 7 | 5 | 57 | 95.36 | 1.54 | 10.27 |
| 8 | 3 | 59 | 95.25 | 1.54 | 6.19 |
| 9 | 5 | 90 | 94.07 | 1.05 | 9.2 |
| 10 | 3 | 51 | 92.03 | 1.7 | 8.8 |
| 11 | 4 | 49 | 91.83 | 1.73 | 18.55 |
| 12 | 2 | 49 | 90.44 | 1.77 | 11.9 |
| 13 | 3 | 54 | 89.03 | 1.56 | 7.08 |
| 14 | 5 | 42 | 82.16 | 1.75 | 8.45 |
| 15 | 1 | 41 | 80.26 | 1.91 | 11.1 |
| 16 | 5 | 39 | 75.3 | 1.71 | 8.38 |
| 17 | 3 | 59 | 74.67 | 1.2 | 9.63 |
| 18 | 3 | 44 | 74.56 | 1.59 | 8.03 |
| 19 | 2 | 30 | 74.29 | 2.32 | 9.77 |
| 20 | 2 | 28 | 71.28 | 2.38 | 11.73 |
| 21 | 3 | 10 | 69.87 | 5.37 | 21.81 |
| 22 | 0 | 29 | 63.29 | 2.18 | 6.86 |
| 23 | 3 | 42 | 53.93 | 1.2 | 8.07 |
| Total | 75 | 1126 | 2008.87 | 1.67 | 9.93 |

### Addition of segregation distorted markers to the linkage map

To localize the significantly segregation distorted (SD) loci within the map, all of the 902 SD markers were tested for linkage group assignments relative to positions of the previously mapped 1,621 non-distorted markers. As a result, 389 SD markers were mapped to 257 distinct locations within the 23 LGs. Most of the SD markers (74%) were observed on linkage groups 1, 2, 5, 8, 13, 18, 19, 21 and 23 (Fig.3, Table S2). Additionally, we observed that LG21 in the maternal map could be generated only when the SD markers were included.

**Figure 3:**
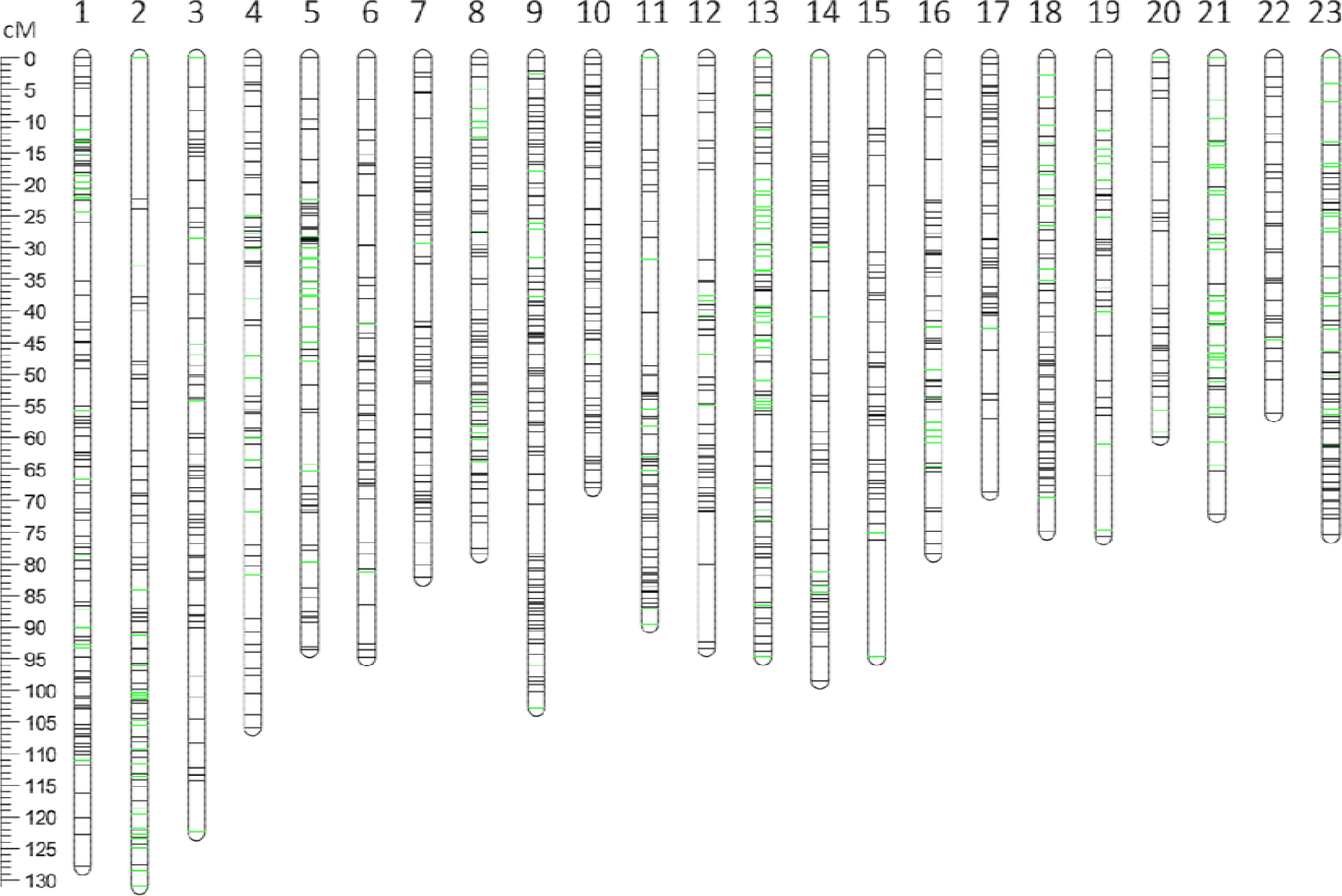
Genetic maps of *F. pennsylvanica* including segregation distorted markers. Green bars represented the segregation distorted loci.

### Comparative analysis of green ash with other species

Tomato (*Solanum lycopersicum*, Sl, 2n=24), coffee (*Coffea canephora*, Cc, 2n=22) and green ash (Fp) belong to the sister orders Solanales, Gentianales and Lamiales, respectively within the asterids. Peach (*Prunus persica*, Pp, 2n=16) and poplar (*Populus trichocarpa*, Pt, 2n=38) are two species from the rosids, which are distantly related to green ash. The 50bp flanking sequences of both sides of the mapped 1,522 SNP markers (1,126 unique locations) were aligned to the genomes of the four species using BLASTN analysis, with an e-value cutoff of 1e-5, revealing that 325 (21.35%), 342 (22.47%), 329 (21.62%) and 239 (15.7%) SNP markers could be mapped to the chromosomes of tomato, coffee, poplar and peach, respectively. The extent of syntenies of the green ash genetic map to these four species was summarized in Tables 5-8. When the queried LG marker sequences and target chromosome genome sequences shared at least 5 loci, the region was considered as a potential syntenic block in our study. For the green ash – tomato comparison, this resulted in 13 syntenic blocks involving 90 loci (28% of shared loci) on 10 ash LGs. For the green ash – coffee comparison, this resulted in 21 syntenic blocks involving 52 loci (15% of shared loci) on 15 ash LGs. For the green ash – peach comparison, this resulted in 12 syntenic blocks involving 83 loci (35% of shared loci) on 6 ash LGs. For the green ash – poplar comparison, this resulted in 9 syntenic blocks involving 49 loci (15% of shared loci) on 10 ash LGs. We constructed circular plots of potential syntenic blocks between each Fp LG and the pseudomolecules of relevant Sl, Cc, Pp and Pt chromosomes (Fig. 4).

**Table 5.**
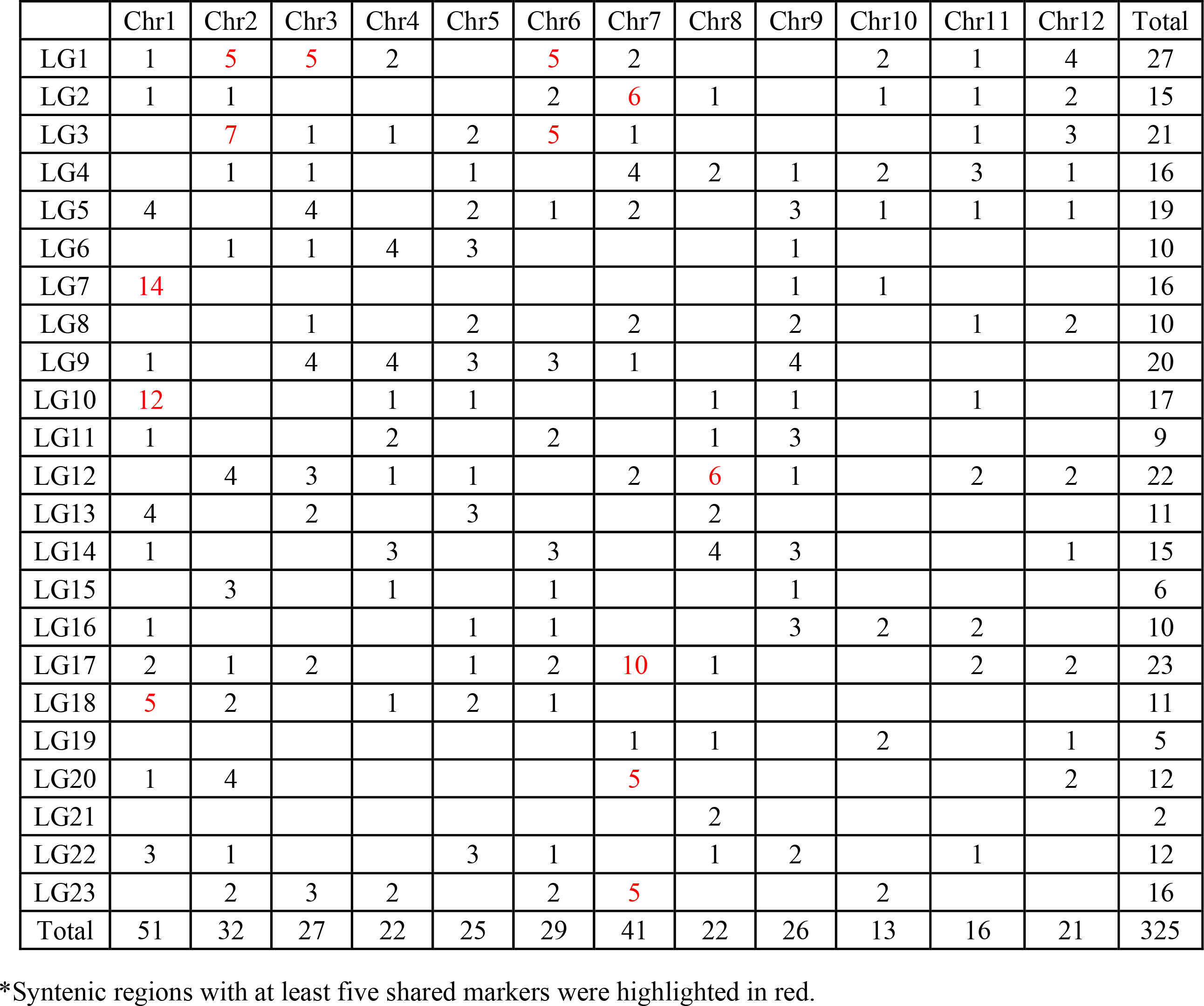
Distribution of orthologous loci on LGs of green ash and tomato genome*.

**Table 6.** Distribution of orthologous loci on LGs of green ash and coffee genome*.

|  | Chr1 | Chr2 | Chr3 | Chr4 | Chr5 | Chr6 | Chr7 | Chr8 | Chr9 | Chr10 | Chr11 | Total |
| --- | --- | --- | --- | --- | --- | --- | --- | --- | --- | --- | --- | --- |
| LG1 |  | 1 | 1 | 9 |  | 2 | 5 | 1 |  | 6 |  | 25 |
| LG2 | 3 |  |  |  | 7 |  |  | 3 |  | 1 | 1 | 15 |
| LG3 | 1 | 1 |  | 7 |  | 1 | 8 | 1 |  | 1 |  | 20 |
| LG4 |  | 2 |  | 1 | 2 | 5 |  | 3 | 2 |  |  | 15 |
| LG5 | 3 | 4 | 1 | 1 |  | 7 |  | 1 | 1 | 1 |  | 19 |
| LG6 |  |  |  |  |  |  | 5 |  |  | 4 |  | 9 |
| LG7 | 3 | 14 |  |  |  |  |  |  |  |  |  | 17 |
| LG8 | 1 | 3 | 1 |  |  | 7 | 2 |  |  | 1 | 1 | 16 |
| LG9 | 8 | 4 | 3 | 5 |  |  |  | 1 | 1 | 2 | 7 | 31 |
| LG10 |  | 14 | 1 |  |  |  |  | 1 |  |  |  | 16 |
| LG11 | 3 |  | 3 |  |  |  |  | 2 |  | 1 |  | 9 |
| LG12 |  | 2 | 2 |  | 3 |  | 1 | 7 | 3 |  | 1 | 19 |
| LG13 | 2 | 1 |  | 2 |  | 1 |  | 2 |  | 1 | 4 | 13 |
| LG14 | 5 | 1 | 3 |  |  |  | 1 | 4 |  | 4 |  | 18 |
| LG15 |  | 1 | 1 |  |  |  | 1 |  |  | 2 |  | 5 |
| LG16 | 5 |  | 1 |  |  | 2 |  |  |  |  | 1 | 9 |
| LG17 | 2 | 1 | 2 | 4 | 1 | 9 | 5 |  | 1 |  |  | 25 |
| LG18 | 1 | 2 |  |  | 2 | 1 |  | 1 | 3 |  | 2 | 12 |
| LG19 |  |  |  | 1 |  | 7 |  |  |  |  |  | 8 |
| LG20 |  | 2 |  | 1 | 3 |  |  |  | 3 |  |  | 9 |
| LG21 |  |  |  |  |  |  |  | 2 |  |  |  | 2 |
| LG22 | 2 | 4 |  | 1 |  |  |  | 2 | 2 |  | 1 | 12 |
| LG23 | 1 | 3 | 1 | 4 | 4 | 1 |  |  |  | 3 | 1 | 18 |
| Total | 40 | 60 | 20 | 36 | 22 | 43 | 28 | 31 | 16 | 27 | 19 | 342 |

\*Syntenic regions with at least five shared markers were highlighted in red.

**Figure 4:**
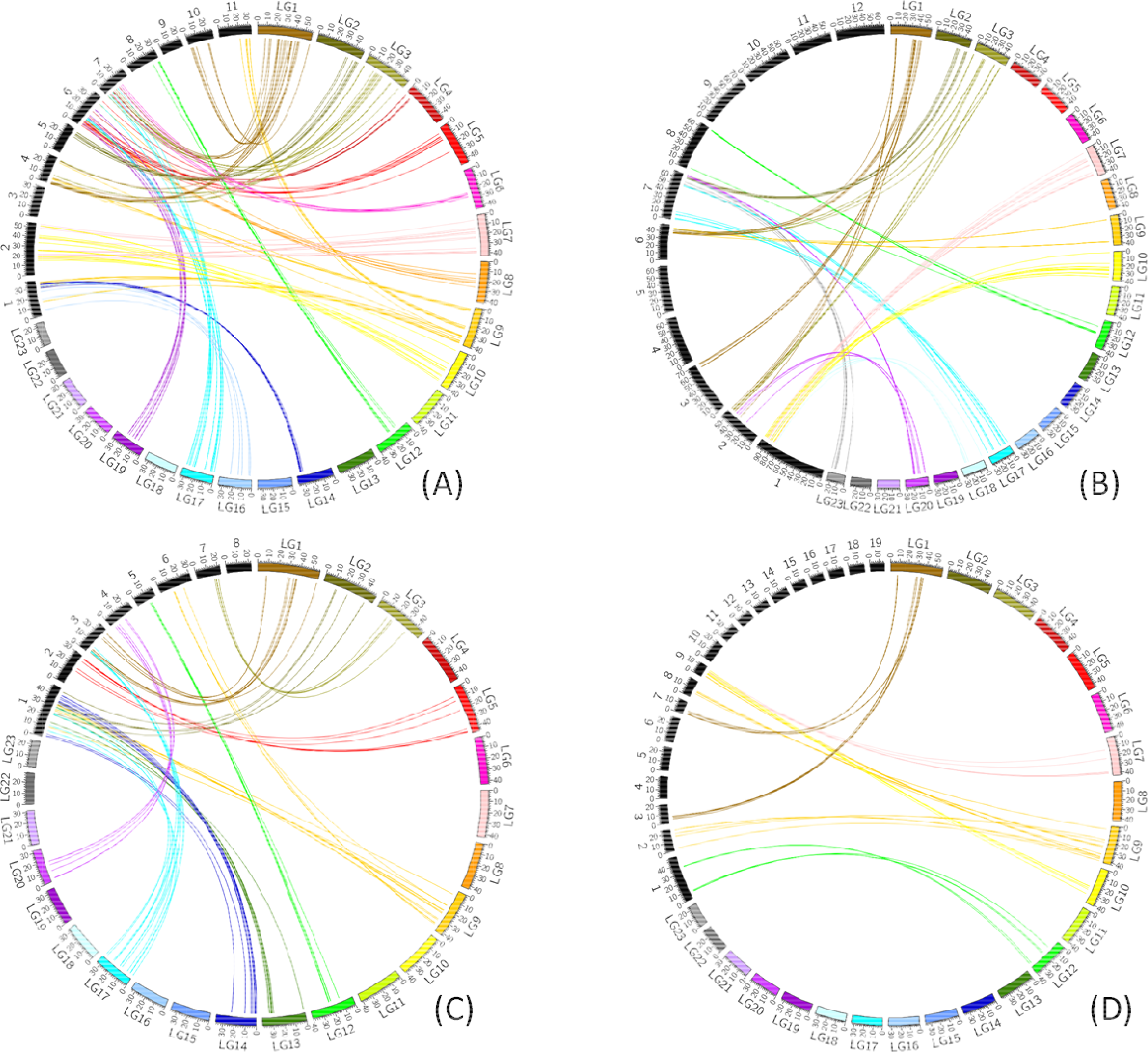
Syntenic relations between the 23 *F. pennsylvanica* linkage groups and each of the 12 tomato chromosomes (A), each of the 11 coffee chromosomes (B), each of the 8 peach chromosomes (C), each of the 19 poplar chromosomes (D), using GBS-derived SNPs. Green ash linkage groups were color coded and labeled from LG1 to LG23. Pseudomolecules of the four target species were shown in black. Links connected the locations of homologs between genomes.

We observed that the order of the syntenic block relationships were inconsistent across the 23 Fp LGs. For example, LG1 of green ash showed unique one-to-one correspondence with the peach genome, and showed a one-to-two correspondence with the poplar genome, but showed a one-to-three correspondence with both the tomato and coffee genomes. As another example, green ash LG12 showed unique one-to-one correspondence with all four target genomes. On the other hand, LG17 showed better conserved synteny with tomato as it corresponded with one chromosome of tomato (Sl_Chr7), but two chromosomes of both coffee (Cc_Chr6 and 7) and peach (Pp_Chr1 and 3). LG2, 13 and 14 of green ash showed syntenic regions with Pp_Chr1; while LG5, 1, 20, 22 and 3 corresponded to Pp_Chr2, 3, 4, 5 and 7, respectively (Table 7). Loci from LG9 were mapped to Pp_Chr1 and 6, and loci from LG17 were mapped to Pp_Chr1 and 3. Synteny of green ash linkage groups with the poplar genome is more eroded than the synteny of green ash with genomes of peach, coffee and tomato. LG7 and 10 suggested a syntenic block with Pt_Chr9. LG12 and 20 showed syntenic blocks with Pt_Chr1 and 11, respectively. Other linkage groups from green ash showed synteny with several additional chromosomes of poplar.

**Table 7.** Distribution of orthologous loci on LGs of green ash and peach genome*.

|  | Chr1 | Chr2 | Chr3 | Chr4 | Chr5 | Chr6 | Chr7 | Chr8 | Total |
| --- | --- | --- | --- | --- | --- | --- | --- | --- | --- |
| LG1 | 4 | 1 | 10 | 2 | 1 | 2 | 3 |  | 23 |
| LG2 | 5 |  | 3 | 4 | 1 |  |  |  | 13 |
| LG3 | 3 |  | 4 | 1 |  | 2 | 5 |  | 15 |
| LG4 |  | 1 | 3 | 2 | 1 | 1 | 3 |  | 11 |
| LG5 | 2 | 7 |  |  |  | 2 | 1 | 3 | 15 |
| LG6 |  |  | 1 |  |  | 1 |  |  | 2 |
| LG7 | 4 | 3 |  |  |  | 1 |  | 3 | 11 |
| LG8 | 2 |  | 2 |  |  |  | 2 |  | 6 |
| LG9 | 9 | 1 |  |  | 2 | 6 |  | 2 | 20 |
| LG10 | 1 | 2 | 1 |  | 1 | 2 | 4 | 3 | 14 |
| LG11 | 4 |  |  |  |  | 1 | 1 | 1 | 7 |
| LG12 |  | 3 |  | 4 | 5 |  | 1 | 1 | 14 |
| LG13 | 7 |  |  |  | 1 | 3 |  |  | 11 |
| LG14 | 9 |  |  |  |  | 2 |  | 1 | 12 |
| LG15 | 2 |  |  |  |  | 1 | 1 |  | 4 |
| LG16 |  |  |  | 2 |  | 4 |  |  | 6 |
| LG17 | 5 |  | 7 |  | 1 | 2 | 3 |  | 18 |
| LG18 | 1 | 1 | 2 |  |  | 2 | 3 | 1 | 10 |
| LG19 | 1 | 1 |  |  |  |  | 2 |  | 4 |
| LG20 |  | 1 |  | 8 |  |  |  |  | 9 |
| LG21 |  |  |  |  |  |  |  |  | 0 |
| LG22 |  | 1 |  |  |  | 1 | 1 | 2 | 5 |
| LG23 |  |  | 2 | 3 | 3 |  |  | 1 | 9 |
| Total | 59 | 22 | 35 | 26 | 16 | 33 | 30 | 18 | 239 |

\*Syntenic regions with at least five shared markers were highlighted in red.

**Table 8.** Distribution of orthologous loci on LGs of green ash and poplar genome.

|  | Chr<br>1 | Chr<br>2 | Chr<br>3 | Chr<br>4 | Chr<br>5 | Chr<br>6 | Chr<br>7 | Chr<br>8 | Chr<br>9 | Chr<br>10 | Chr<br>11 | Chr<br>12 | Chr<br>13 | Chr<br>14 | Chr<br>15 | Chr<br>16 | Chr<br>17 | Chr<br>18 | Chr<br>19 | Total |
| --- | --- | --- | --- | --- | --- | --- | --- | --- | --- | --- | --- | --- | --- | --- | --- | --- | --- | --- | --- | --- |
| LG1 | 1 |  | 5 |  | 4 | 1 | 5 | 2 | 1 | 2 | 1 | 3 |  | 2 |  |  | 2 |  |  | 29 |
| LG2 | 2 |  | 1 |  |  |  |  | 1 |  | 1 | 2 |  | 1 |  |  |  |  | 3 | 2 | 13 |
| LG3 | 3 |  | 3 |  | 1 | 1 | 4 |  |  | 1 |  | 2 |  | 2 | 1 |  | 2 | 1 |  | 21 |
| LG4 | 1 |  | 1 | 2 |  | 1 |  |  |  |  | 2 |  | 2 | 1 |  |  |  | 4 |  | 14 |
| LG5 |  | 4 | 1 | 1 |  | 3 |  |  | 1 | 1 |  | 3 | 1 |  | 1 | 1 | 2 | 2 | 1 | 22 |
| LG6 |  | 2 |  |  | 4 |  |  |  |  |  |  |  |  |  |  |  | 1 |  | 1 | 8 |
| LG7 |  |  | 2 | 1 |  |  |  | 4 | 6 |  |  |  |  | 1 |  |  |  |  |  | 14 |
| LG8 | 1 |  |  |  | 1 | 3 | 1 |  |  | 4 |  |  | 1 |  |  |  |  | 2 |  | 13 |
| LG9 | 3 | 5 |  |  | 2 |  | 1 | 5 | 5 | 1 |  | 3 |  |  | 3 |  |  |  |  | 28 |
| LG10 | 1 |  |  | 2 | 1 |  |  |  | 6 |  |  |  | 2 | 3 | 1 |  | 1 |  |  | 17 |
| LG11 | 1 |  | 3 |  |  |  |  |  |  |  |  |  |  | 2 |  |  |  |  |  | 6 |
| LG12 | 6 |  | 2 |  | 1 |  |  |  |  | 2 |  | 3 |  |  |  |  |  |  | 1 | 15 |
| LG13 | 4 | 1 | 1 |  |  |  |  |  |  | 3 |  |  |  | 1 | 2 | 1 |  | 2 |  | 15 |
| LG14 | 4 | 3 | 1 |  | 2 | 3 |  |  |  | 1 |  |  | 1 | 3 |  | 1 |  | 2 |  | 21 |
| LG15 | 1 | 1 |  | 1 | 1 | 2 |  |  |  | 1 |  |  |  | 1 |  |  | 1 |  |  | 9 |
| LG16 | 3 |  |  |  |  | 1 |  |  | 1 |  |  |  |  | 1 |  | 1 |  |  |  | 7 |
| LG17 | 2 |  | 3 | 1 | 2 | 4 |  | 2 | 2 | 1 |  | 1 | 1 | 1 |  |  |  | 4 |  | 24 |
| LG18 |  | 4 |  | 2 |  |  |  | 1 | 2 | 1 |  |  |  |  |  |  |  |  |  | 10 |
| LG19 |  |  | 1 |  |  | 3 |  |  |  |  |  |  |  |  |  | 1 |  | 2 |  | 7 |
| LG20 | 2 |  |  | 1 |  |  |  |  |  | 2 | 6 |  |  | 2 | 1 |  |  |  |  | 14 |
| LG21 |  |  |  |  |  | 1 |  |  |  |  |  |  |  |  |  |  |  |  |  | 1 |
| LG22 |  | 1 | 1 | 3 |  |  |  |  | 1 | 1 |  |  |  | 1 | 1 |  |  |  |  | 9 |
| LG23 | 1 |  |  | 1 |  | 1 | 1 |  |  |  |  | 3 |  |  | 2 |  |  |  |  | 9 |
| Total | 36 | 21 | 25 | 15 | 19 | 24 | 12 | 15 | 25 | 22 | 11 | 18 | 9 | 21 | 12 | 5 | 9 | 22 | 5 | 326 |

Overall, the green ash map showed higher synteny with the coffee genome than the other genomes. For the 23 Fp linkage groups, 11 linkage groups (LG2, 4~8, 10, 12, 14, 16 and 19) showed syntenic regions to a single chromosome of coffee, while 8 linkage groups (LG2, 7, 10, 12, 17, 18, 20 and 23) showed syntenic blocks to a single chromosome of tomato. Compared with rosid species, we found that 8 LGs and 4 LGs showed a one-to-one correspondence with the peach and the poplar genomes, respectively. In addition, 5 green ash LGs only showed syntenic blocks with the coffee genome, and not with any of the other species.

## Discussion

Development of the GBS technology has made it possible to rapidly obtain genotyping data for thousands of loci in months. Restriction enzyme selection is a critical step to target sites in low-copy genomic regions, minimizing reads in repetitive sequences, which influences both the number and genomic location of SNPs discovered. We tested *ApeKI* (GCWGC), *PstI* (CTGCAG) and *Eco*T22I (ATGCAT) enzymes, previously used in maize, pine and other plant species. There is a tradeoff between the read coverage of SNPs and the number of SNPs. For our study, we aimed to genotype at good coverage with at least several hundreds of loci. Hence, we chose the six-cutter enzyme *PstI*, to create reduced representation libraries that produced fewer repetitive fragments and a large proportion of fragments within sequencing range of <500bp. As a result, we discovered 63,540 SNPs, which were mapped to 3,973 reference genome scaffolds. However, only 5727 (9%) of these SNPs exhibited less than 20% missing data. This large proportion of missing data may have resulted from high heterozygosity levels in this out-crossed species. In future studies, the missing data could be lowered to reduce the level of multiplex or by constructing optimized two-enzyme libraries.

Our SNP identification revealed a Ts/Tv ratio of 1.78. This level of transition bias has also been observed in many other plant species. Transitions are favored due to their better tolerance than transversions during natural selection, as they are more likely to contribute synonymous mutations in protein-coding regions. A Ts/Tv ratio of 2.00 was detected in selected inbred lines of *Indica* rice (Subbaiyan et al. 2012), while in oil palm, SNPs showed a Ts/Tv ratio of 1.67, using a modified two-enzyme GBS protocol (Pootakham et al. 2015). The ratio between transition and transversion was 1.42 in sunflower (Celik et al. 2016). In chickpea, a Ts/Tv ratio of 1.74 was reported from SNP discovery (Gaur et al. 2015).

Although SNPs can be obtained in higher numbers than SSRs, bi-allelic SNP markers are often less informative than SSRs. Multi-allelic SSR markers are more useful when integrating parental genomic data to create an intra-specific map. SSR markers are also more transferrable among different populations and species. Therefore, a combination of SNPs and SSRs represents an ideal strategy for map construction. SSRs have limitations in terms of cost and time-consumption. We only identified 84 reliable polymorphic SSR markers from the hundreds of SSRs discovered.

Many genetic maps have been reported for some tree species, including *Eucalyptus* sp. (Freeman et al. 2006), *Populus* sp. (Cervera et al. 2001), *Quercus* sp. (Barreneche et al. 1998) and *Pinus* sp. (Neves et al. 2014). However, a genetic map of *Fraxinus pennsylvanica*, a species critically endangered by the invasive emerald ash borer (IUCN red list: Westwood et al. 2017) has not been developed until now. We found that a 96-plex GBS protocol worked well for the construction of a high-density linkage map with a compact marker interval (< 2cM) in a non-model, outcrossing species. Our green ash linkage map was constructed with 1201 markers, including 1126 SNP and 75 SSR loci, spanning 2008.87 cM. The map consisted of 23 linkage groups, equal to the chromosome number, that range from 53.93 to 127.05 cM. A higher recombination rate was observed in the male parent (1.94cM/Mb) versus the female parent (1.74cM/Mb), which was consistent with observations in *Arabidopsis* (Giraut et al. 2011). Among 23 linkage groups, the most dramatic differences in map length between two parents were observed between LG16 and LG18, suggesting that sex-specific patterns may be associated with these two chromosomes. The average recombination rate across all linkage groups was 2.23cM/Mb, which is comparable to rates reported for cacao (1.7cM/Mb), grape (2.0cM/Mb), papaya (2.54cM/Mb) and soybean (2.51cM/Mb) (Henderson 2012).

Segregation distortion (SD) is a general phenomenon in plants, but the percentage, degree and genetic effects may vary significantly across species (Dai et al. 2016; Zhou et al. 2015; Taylor and Ingvarsson 2003). In maize, it has been reported that 18 chromosome regions on 10 chromosomes were associated with SD (Lu, Romero-Severson, and Bernardo 2002) while 14 SD regions were reported for barley (Li et al. 2010). SD has been suggested as a selection mechanism (Sandler and Novitski 1957) and it may result from biological and environmental factors, such as chromosome loss (Bradshaw and Stettler 1994), or gametic and zygotic selection (Liebhard et al. 2003). In our study, 33.02% of the markers showed distortion from the expected segregation ratio. In a future study, with increased population size, we may be able to detect different regions associated with SD markers compared to what we reported here. Unspecified viral infection killed ~350 highly susceptible genotypes in the mapping population, which may have led to segregation distorted markers around the genomic region or regions responding to the viral infection.

Synteny was retained during genome evolution and duplication events. Genome investigations have shown evidence for whole genome duplication (WGD) across all flowering plant lineages. The model organism, *Arabidopsis thaliana*, underwent at least three ancient genome duplication events (α, β and γ) over the last 300 million years (Bowers et al. 2003). Both α and β events occurred within rosids clade II: the α event, shared within genus *Brassica*, and the β event occurred within the order of Brassicales following the divergence from papaya (Bowers et al. 2003). A whole genome triplication, called γ, was suggested by the analyses of sequenced genomes of poplar (Tuskan et al. 2006), grape (Jaillon et al. 2007) and papaya (Ming et al. 2008), which may have occurred close to the eudicot divergence (Soltis et al. 2009). Furthermore, a more recent additional WGD event also occurred in some species, such as poplar (Tuskan et al. 2006), cotton (Wang et al. 2012), genus Brassica (Lysak et al. 2005; Lukens et al. 2004), and the Cleomaceae (Schranz and Mitchell-Olds 2006), while a recent genome triplication occurred in the Solanum lineage (The Tomato Genome 2012).

To understand syntenic relationships between green ash and related species, tomato, coffee, peach and poplar were selected to conduct comparative analyses. Tomato (Solanales) and coffee (Gentianales) are sister orders to green ash (Lamiales) within the Lamiids major clade in the Euasterids. These three orders diverged from their last common ancestor approximately 78-80 (Magallon, et_al. 2009) or 82-89 million years ago (Wikström, Savolainen, and Chase 2001). Only 6 of the 12 tomato pseudomolecules and 8 of the 11 coffee pseudomolecules contained syntenic blocks with the green as map of ≥5 loci. Ten of the 23 green ash LGs contained syntenic blocks with tomato pseudomolecules, while 15 green ash LGs contained syntenic blocks with the coffee pseudomolecules, ranging from 1 to 3 syntenic blocks per LG. These synteny features indicate that genome duplication and rearrangements may have occurred since the separation of the three species, as previously reported for species in all 3 of the orders (Ren et al 2018), including a genome triplication in the tomato genome (Tomato Genome Consortium 2012) that was not found in the coffee genome which does alternately include several species-specific gene family expansions (Denoeud et al 2014). Previous synteny analysis among woody tree species and herbs has suggested generally slower rates of synteny erosion during genome divergence for woody perennials (Luo et al. 2015). Here, we also observed that syntenic relationships between the green ash and coffee woody species were somewhat more conserved than between green ash and the herbaceous tomato, in terms of chromosomes and linkage groups carrying conserved microsyntenic regions, even though phylogenetic distances among the 3 species are equivalent. Poplar and peach are much far more distantly-related to green ash, providing examples of woody species in the two major fabids and malvids groups within the eurosids (Cantino et al. 2007). Even though the Rosid and Asterid clades diverged 125 million years ago (Wikström, Savolainen, and Chase 2001), we still detected micro-syntenic regions between these distantly related woody species and green ash, totaling 9 blocks between 6 green ash LGs and 7 of the 19 poplar chromosomes, and 12 syntenic blocks observed between 10 green ash LGs and 7 of the 8 peach chromosomes. Our results suggest that coffee and peach should be used as model species for further comparative genomic analyses among woody species.

This research allowed us to provide detailed genetic marker data and construct the first reference map for *F. pennsylvanica*. These genetic resources provide a platform for identifying QTLs for traits of importance in future ash breeding programs. The DNA markers identified in this study also can be utilized for analyses of genetic variation and population structure for this threatened ash species, providing valuable information that can guide efforts to conserve genetic diversity for the species in the future. No phenotypic traits have yet been characterized within the present set of progeny in our mapping population as they were still at the seedling stage during construction of the map. The entire mapping family has recently been planted at a site that should expose the progeny to EAB attack. Over the next several years, phenotypic data will be collected for QTL identification, including traits associated with EAB resistance, growth, and budburst. In addition, this genetic map will be utilized to map candidate genes identified from previous expression profiling analyses. The results from this study can be extended to future association mapping analysis as well. Hopefully, the results reported here will facilitate the development of further genetic and genomic tools to identify EAB resistant seedlings for use in tree breeding efforts in the US, ultimately leading to the restoration of resistant green ash trees to support recovery of this important tree species across its native range.

